# Dominant resistance alleles accelerate population suppression by CRISPR-based gene drive

**DOI:** 10.64898/2026.05.28.728369

**Authors:** Weizhe Chen, Jingyun Liang, Jackson Champer

## Abstract

CRISPR-based gene drive is a promising strategy for pest control. However, when targeting a recessive fertility gene, nonfunctional resistance alleles generated by end-joining slow the spread of drive and can even impair suppressive power. It was recently discovered that a *doublesex* target site can result in dominant female-sterile alleles when disrupted. It could serve as an ideal target for suppression drives, as disrupted alleles are efficiently removed when inherited by a female, increasing the suppressive power of the drive. Here, to better understand how dominant sterile target sites can improve suppression, we used an individual-based, continuous space model to investigate chasing dynamics of homing drives that produce dominant female-sterile resistance. Our results indicate that dominant-sterile resistance allows homing drives to achieve higher population elimination rates and a lower probability of chasing over a broader range of performance parameters. In addition, we propose a confined gene drive system, termed TADFI (Toxin-Antidote Dominant Female Intersex), that is also based on formation of dominant female sterile alleles. TADFI has two variants, designed for population suppression and modification, respectively. We estimate the introduction frequency threshold of TADFI in a panmictic model, and evaluate its suppression dynamics in a spatial model. Though spreading less rapidly than Toxin-Antidote Dominant Embryo (TADE) suppression drive, it has similar favorable dynamics related to confinement and suppressive power, while potentially being easier to construct. Together these results show that drives generating dominant-female sterile alleles may be promising candidates for a variety of population suppression applications.

## Introduction

Pest insects impose severe burdens on agriculture, the economy, and public health worldwide. Many species act as vectors of human and animal diseases^1–3^. Others are responsible for substantial crop losses, reduced food security, and billions of dollars in economic damage every year^1,4,5^. Current control strategies primarily rely on chemical insecticides, biological control agents, and sterile insect technique (SIT)^6–8^. However, these approaches are often constrained by the rapid evolution of resistance, high costs of large-scale implementation, and limited long-term sustainability^9,10^.

Gene drives are selfish genetic elements that bias their own inheritance, allowing them to be transmitted to offspring at super-Mendelian frequencies (>50%)^11^, They can be designed to spread desired traits throughout wild populations or directly eliminate target populations. CRISPR techniques have allowed more flexible and practical gene drive designs, boosting efficiency and offering high potential for future pest control. CRISPR-based gene drives are mainly divided into homing drives, which propagate by copying themselves into target sites, and toxin-antidote drives, which act by killing non-drive individuals while rescuing drive carriers^12^. Together, these strategies have been utilized to developed systems for population suppression or modification in mosquitoes^13–16^, flies^17–21^, *Arabidopsis*^22,23^, mice^24^ and moths^25^, though no gene drive has yet been deployed in the field.

Homing drives employ CRISPR-mediated cleavage and homology-directed repair to convert heterozygotes into homozygotes in germline cells, thereby biasing their inheritance. However, double-strand breaks can also be repaired through end-joining pathways, which may generate resistance alleles that are no longer recognized by Cas9, consequently slowing down or even completely blocking population suppression^18,26,27^. Haplosufficient genes, including *doublesex*^14–16,19,28^, *yellow-g*^20^, *transformer*^21^, and *sxl*^21^, are frequently selected as targets in homing suppression drives. Female homozygotes remain fertile and able to pass on the drive to most offspring. When the drive reaches high frequency, there are many sterile female homozygotes, causing population collapse. By using multiple gRNAs and targeting conserved regions of genes, the risk of functional resistance can be mitigated^14,20,29,30^. However, formation of nonfunctional resistance alleles can still reduce the suppressive power (genetic load) of the drive, potentially preventing complete population elimination, even if the drive remains at high equilibrium frequency^20,31^. Targeting the female-specific splicing acceptor site within *doublesex* can yield dominant female sterile resistance alleles with the drive itself still being recessive sterile^18,28,32^. This allows stronger suppressive power compared with the drives that produce only recessive resistance alleles^18^. Specifically, formation of dominant female-sterile alleles actually increases genetic load, even if the drive has fitness costs.

Previous studies showed that deployment of suppression drives may have additional complexities in spatially explicit populations^33–40^. One of the most prominent is the “chasing” effect, in which some wild-type individuals elude the drive and recolonize previously suppressed areas. Chasing population dynamics may postpone target population elimination or even cause the drive to fail ^41–45^. The chasing phenomenon frequently occurs when the drive exhibits low performance, even if performance is still high enough to avoid an equilibrium state and succeed in a panmictic model^33,40,46^. Dominant sterile resistance homing suppression drives can also have low drive performance, especially if drive homozygous males are also sterile, as reported experimentally^18^. It remains unclear whether such a drive would perform well in spatial environment. Here, we compared in spatial models several types of homing suppression drives that (i) generate recessive female sterile resistance alleles, (ii) generate dominant female sterile resistance alleles, and (iii) generate dominant female sterile resistance alleles but with an extra effect of producing sterile drive homozygous males. We showed that a target site producing dominant female-sterile alleles accelerates population elimination across a broader range of parameters in spatial models, even when the homozygous males were sterile.

Apart from homing drives, another type of CRISPR-based gene drive system, called toxin-antidote drives, increase drive allele frequency by targeting and disrupting wild-type alleles (the “toxin”), which are then removed from the population. Toxin-antidote drives have protective rescue element (the “antidote”), ensuring that individuals carrying the drive can usually survive and reproduce^47,48^. Toxin-antidote drives have the advantage of being confined to target populations due to their frequency-dependent dynamics. In their idealized form, basic forms of toxin-antidote drives typically lack an introduction threshold (the frequency below which the drive will fail). However, embryo cleavage caused by deposited Cas9 (in drives that create dominant-effect disrupted alleles) or fitness costs can impose a threshold, thus resulting in confinement. Some underdominance drive forms also have thresholds even with ideal performance^49^. Most variants can only be applied to population modification, but some can also be adapted for suppression, such as the Toxin-Antidote Dominant Embryo (TADE) suppression drive^46,48^. TADE drives disrupt a haplolethal gene through CRISPR/Cas9-mediated cleavage, so resistance alleles are dominant lethal, resulting in their immediate removal from the population^46,48,50^. To achieve suppression, the drive is inserted into a haplosufficient female fertility gene, such that homozygous females carrying the drive are sterile (alternatively, they can be elsewhere and target the female fertility gene with additional gRNAs). While promising, including in spatial models^46,51^, TADE-suppression transgenic lines are challenging to establish^52^, and there has thus far not been a successful demonstration.

Here, we propose a variant named TADFI (Toxin-Antidote Dominant Female Intersex), instead of targeting a haplolethal gene, it targets a female-specific locus that produces dominant sterile disrupted alleles, such as the site in *doublesex*^28,32^. This gives it the similar threshold-dependent dynamics, and it could be similarly converted to a suppression drive. Although the efficiency of TADFI would be weaker than that of TADE, it is can potentially be developed more easily because the disrupted alleles do not affect males and rescue element expression need to be as finely regulated. Considering chasing dynamics of TADE suppression systems in a previous study^46^, our newly proposed threshold-dependent TADFI systems are likely to be more vulnerable to chasing due to their reduced ability to spread. To better predict and understand the key features of how this drive behaves, we evaluated its characteristics and performance in both panmictic and spatial models. As long as embryo cutting remains at a low level and the germline cut rate is at least moderate, population elimination can be achieved even in the presence of chasing.

## Methods

### Drive mechanisms in this study

Our model includes wild-type alleles, drive alleles, and (nonfunctional) resistance alleles. For the HSD-recessive system, both the drive allele and resistance allele can induce recessive female sterility.

For the HSD-dominant system, the drive allele causes recessive female sterility, whereas the resistance allele leads to dominant female sterility. In one variant of the HSD-dominant system, the drive allele also results in recessive male sterility (resistance alleles do not affect male fertility). Sterile males do not participate in mating (they are intersex). All homing drives allow drive/wild-type heterozygous individuals to convert a fraction of wild-type alleles in their germline into drive alleles at the drive conversion rate. Wild-type alleles can also be converted to resistance alleles at the germline resistance formation rate. In some simulations, a proportion of resistance alleles can be functional. These are equivalent to wild-type alleles, except that they cannot be affected by the drive. Fitness costs are considered for drive heterozygous female in these systems, usually representing the effect of undesired Cas9 activity, such as in somatic cells. Fitness affects female fecundity. Note that drive/functional resistance allele heterozygotes suffer no female fitness costs.

In the TADE suppression system, the drive element need not placed at a locus close to the target site, but in this study, they are assumed to be closely linked (to facilitate direct comparisons with TADFI, though this is not expected to affect performance for an ideal drive). All disrupted alleles (equivalent to nonfunctional resistance alleles for CRISPR toxin-antidote systems) exert a dominant lethal effect at the embryo stage unless the zygote carries sufficient copies of the drive allele (no less than the number of disrupted alleles) to provide rescue^46,48^.

In the TADFI system, the drive and the target site are located at the same locus, a haplosufficient female fertility gene, but the disrupted alleles are dominant sterile. Drive/wild-type heterozygous individuals can disrupt a fraction of wild-type alleles in their germline at the germline resistance formation rate. For the suppression design, no rescue is provided for the drive allele, so drive homozygous females will be sterile. Fitness costs are specifically considered in drive heterozygous females as above. For the modification design, the drive allele carries a rescue element, so all drive homozygotes would be fertile. Fitness costs is applied for each drive allele multiplicatively and affect female fecundity and male mating success.

Considering maternal deposition of Cas9, a fraction of wild-type alleles can be converted into resistance/disrupted alleles at the embryo stage, depending on the embryo resistance formation rate and the maternal genotype^27^.

### Panmictic model

Stochastic simulations were conducted in SLiM (version 4.0)^53^, following approaches similar to those in previous studies^18,28,48^. We modeled a single panmictic population of generic diploids with discrete generations, where the population state was defined by the numbers of male and female adults of each genotype. In each generation, every adult female randomly selected a mate from the adult male pool to produce the next generation. To incorporate crowding and competition effects typical of insect populations, female fecundity was determined by its genotype and total population density. Each offspring was assigned a random sex, and its genotype was determined by randomly inheriting one allele from each parent at every locus, with adjustments for drive activity. Each simulation had a starting population of 100,000 individuals. The population was allowed to equilibrate for 10 generations before introducing gene drive individuals.

### Threshold finder

We employed a threshold finder to precisely determine the introduction frequency threshold of the drive required for successful drive spread under different parameter conditions. The introduction ratio is defined as the fraction of released individuals relative to the total population size immediately after release. The threshold finder began with an introduction ratio of 0.5. If the drive failed (decreasing in frequency toward zero), the ratio was increased by half of the current step size; if the drive succeeded (increasing in allele frequency for a few generations), the ratio was decreased by half of the current step size. The initial step size was set to 0.25 and was iteratively halved in subsequent rounds. This process repeats six times to get an initial estimate of the threshold. To get an exact estimate, we systematically assessed introduced levels in 1% increments starting 0.05 below the initial estimate, performing 10 simulations per value. If the number of failed simulations exceeded 5 (half of total tests), we increased the introduction ratio by 0.01 and tested again. This process was repeated until the number of failed simulations was fewer than 5. The resulting introduction ratio was considered to be the threshold under the current parameter set, representing the boundary between success and failure of this system. If the drive could not succeed when the introduction ratio was set to 0.99, we recorded the threshold as N.A., representing a system that cannot spread under any circumstances.

### Two-dimensional spatial model

In two-dimensional spatial models, females can only sample males within a circle of radius equal to the *average dispersal distance* rather than the whole population.

Female fecundity is determined by local population density. Specifically, it depends on the number of individuals within a circular neighborhood of radius equal to the density interaction distance around the focal female (*ρi*), a carrying density constant (*ρ*, equal to population carrying capacity in total area, set to 100,000 in all simulations except the simulations in Fig.1), the fitness of the female (*f*), the maximum number of offspring per female (*βmax*), and the population growth rate in low density (*β*) (in the absence of competition). The fecundity of given female is defined as follows:

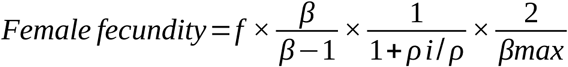

**Figure 1.**
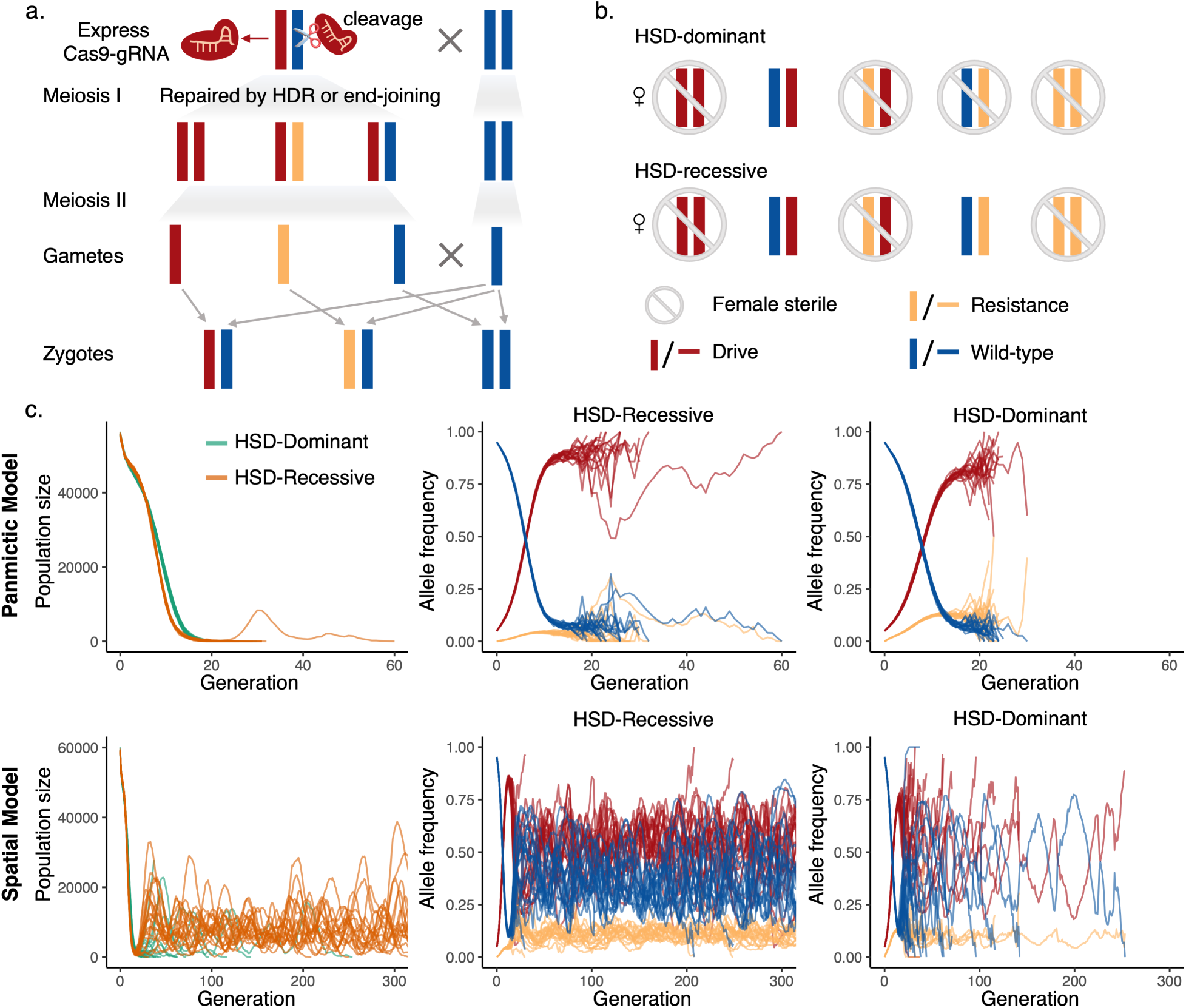
Comparison of HSD-dominant system and HSD-recessive system in panmictic and spatial models. **(a)** Inheritance scheme of CRISPR homing drive. Germline drive conversion, mediated by CRISPR system and HDR, occurs in the germline cells of drive heterozygous parent. Resistance allele formation can also take place. **(b)** Genotypes and sterility in the HSD-dominant and HSD-recessive systems. **(c)** Simulations in a panmictic or spatial populations and with HSD-dominant or HSD-recessive systems. Panels show allele frequencies and populations sizes after an initial release of 10% drive heterozygotes. We parameterized the two systems to produce comparable genetic load (approximately 0.9). For the HSD-dominant system, drive conversion = 0.6, and germline resistance = 0.2. For the HSD-recessive system, drive conversion = 0.8, and germline resistance = 0.0. In both systems, embryo resistance = 0.1, and drive-carrier female fitness at 0.8. Other model parameters are at default levels (Supplementary Table1).

Offspring are displaced from their mother in a random direction at a distance drawn from a normal distribution with a standard deviation equal to the *average dispersal distance*. This produces an average displacement of *average dispersal distance*. If an individual is placed outside the arena, its position is redrawn until it is placed in the arena.

Chasing detection criteria in continuous space is based on a previous method^33^. We quantified spatial clustering at each time step using Green’s coefficient^54^. The landscape was divided into an 8*×*8 square grids. *ni* is the number of wild-type homozygotes in grid *i*, and we denoted the mean and variance of *n_1_* ,…, *n_64_* as *n*and *s*^2^, respectively. N is the number of wild-type homozygotes of all grids. Thus, Green’s coefficient is:

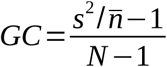

*GC* approaches 0 when individuals are randomly distributed and approaches 1 when they are maximally clustered. A time-series of the inferred *GC*(*t*) and *N*(*t*) (number of wild-type alleles) values was recorded during each simulation run. The drive initially clears the population radially from the release center, causing wild-type individuals to cluster along the edges of the landscape. This leads to an increase in *GC(t)* and a decrease in *N(t)*. However, when wild-type individuals disperse into low-density areas and reproduce, this pattern reverses. We identified the first local maximum in *GC(t)* and the first local minimum in *N(t)*, requiring that each extremum be supported by at least three generations on either side with less extreme values. This indicates the onset of wild-type recolonization and expansion, with increasing wild-type individuals and decreasing clustering. We classified this event as a chase, and the timepoint of the extrema as the chasing start generation.

### One-dimensional spatial model

The one-dimensional space model was similar to two-dimensional model. The arena has a length of 31.33. Individuals were allowed to move and mate within a limited one-dimensional distance (*average dispersal distance,* 0.04), while the density interaction distance was 0.01. The reproduction and local competition were considered within *average dispersal distance*. The one-dimensional model was used for the measurement of drive wave speed and width.

### Drive wave speed measurement

We divided the whole one-dimensional space into 10 equal slices. Drive individuals were released in the leftmost slice, and form an advancing wave toward the right. To calculate the wave speed, we selected the third and eighth slices from the left as reference positions for measurement, which were found to be compatible with this requirement while allowing drive wave measurement over a sufficiently long length for high accuracy.

For homing drives and the TADE suppression system, we used drive allele frequency > 0.5 as a marker for the advancing wave. When the gene drive frequency in the third slice was more than 0.5 for the first time, we defined the generation as the “start generation” and the frequency *Freq_ _Start_* as “start frequency”. The generation before the start generation is defined as *Gen__Start0_*, with a drive frequency of *Freq_ _Start0_*,. Then we calculated the “exact” start generation as:

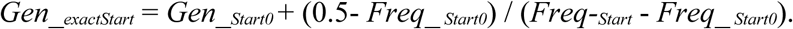

This approximation minimizes the error introduced by discrete generations and is intended to estimate the hypothetical continuous generation time at which the drive frequency reaches 0.5. We then repeated this process for the eighth slice to approximate the exact “stop generation”. When the gene drive frequency in the eighth slice was more than 0.5 for the first time, we defined the generation as “stop generation” and the frequency *Freq__Stop_* as “stop frequency”. The previous generation *Gen__Stop0_* and the its frequency *Freq__Stop0_*were recoded. Thus, the exact stop time is:

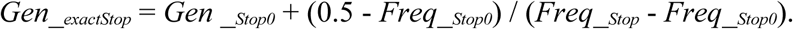

For TADFI suppression system, the advancing equilibrium level of the drive allele frequency was sometimes too close to 0.5, resulting in inaccurate measurements. Thus, we use the state of wild-type allele frequency < 0.8 as the trigger point for measurement. The recorded frequency here is the wild-type allele frequency, and the formulas are changed to:

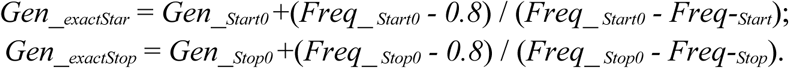

We recorded the time at which the drive wave passed slice 3 and slice 8, with the distance between these two slices corresponding to half of the total arena. The wave speed was therefore calculated as:

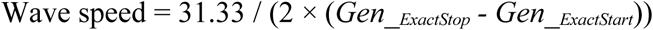

### Wave width measurement

We divided the one-dimensional arena into 50 slices and reported genotype frequencies in each when the wave had reached roughly the middle of the arena. Drive individuals were released in the leftmost 10% region, and form an advancing wave toward the right.

The drive wave width is the distance between the positions where the carrier frequency is 10% and 90%. For each applicable timepoint, we selected the first slice where the drive carrier frequency is lower than 90%. We used the midpoint of each slice as its position. We noted the position of this slice as *pos9.1*, and the drive carrier frequency in this slice as *freq9_1*. The position of the slice that is on the left of *pos9_0* is defined as H0, with drive carrier frequency *freq9_0*. The first slice where the drive carrier frequency is lower than 10%. The position of this slice is marked as *pos1_1*, and the drive carrier frequency in this slice is marked as *freq1_1*. The position of the slice which is to the left of *pos1_1* is marked as *pos1_0*, which has a drive carrier frequency of *R*0. Since the arena at a total length of 31.33, the length of each slice is 0.6266. Thus, the estimated exact positions of start position and stop position are as follows:

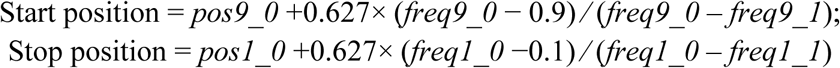

For TADFI suppression drive, since the wave width is relatively board, which decreases the measurement accuracy in the arena of length 31.33, we extended the arena to 62.66 by doubling the starting population size and capacity and halving the competition distance and dispersal rate. We used eligible waves from 20 simulations to find an average wave width.

### Code Availability

Modeling code and all collected data have been deposited on GitHub (https://github.com/chenwz22/TADFI_supplement/).

## Results

### General comparison of suppression drives with dominant and recessive resistance alleles

A previous study demonstrated that dominant female sterile resistance can improve homing suppression drive by increasing genetic load (an indicator of suppressive power)^18^. This occurs because resistance alleles are rapidly eliminated by the dominant sterile effect once transmitted to females, limiting their accumulation in the population while simultaneously contributing to female sterilization. Specifically, dominant resistance can increase the genetic load nearly as well as increased drive conversion, while recessive resistance alleles never increase genetic load and can substantially reduce it in the presence of drive fitness costs. However, these studies were conducted primarily in panmictic models. Outcomes in spatial models, especially for self-sustaining drives, are more complex. In spatially structured populations, once a local region is completely suppressed, the resulting empty space provides an opportunity for wild-type individuals to recolonize the region, a dynamic we refer to as the “chasing” phenomenon^33,42–45^.

To better understand how dominant resistance impacts population dynamics in spatial models, we compared HSD-dominant and HSD-recessive systems. In both systems, the drive allele confers recessive female sterility. However, in the HSD-dominant system, resistance alleles cause dominant female sterility, whereas in the HSD-recessive system, they are recessive female sterile (Fig. 1a). Note that we initially consider only nonfunctional resistance alleles with these female-sterile effects (see below for functional resistance allele considerations). We parameterized the two systems to produce comparable genetic load (approximately 0.9). Specifically, for the HSD-dominant system, drive conversion was set to 0.6 and germline resistance to 0.2. For the HSD-recessive system, drive conversion was set to 0.8 with no germline resistance. In both cases, embryo resistance was fixed at 0.1, and drive-carrier female fitness at 0.8.

In the panmictic model, both the HSD-dominant and HSD-recessive systems were able to suppress the entire population within 40 generations, except for one replicate, which took somewhat longer (Fig. 1b). Despite the lower drive conversion efficiency, the HSD-dominant system was still slightly faster at eliminating the population. In the spatial model, we allowed the simulations to extend up to 1000 generations (Fig. 1c). For the HSD-dominant system, all 20 replicates achieved population elimination within 300 generations, despite transient chasing events. In contrast, the HSD-recessive system exhibited greater variability. Among 20 replicates, 10 continued chasing for 1000 generations, 7 achieved population elimination after chasing (only 3 within 300 generations), and in 3 cases the drive was lost after chasing. These results indicate that while the two systems perform similarly in a panmictic context, the HSD-dominant system substantially outperforms the HSD-recessive system in spatial models, even when suppressive power is the same. One possible explanation is that, in the HSD-recessive system, wild-type alleles partially protect nonfunctional resistance alleles, allowing resistance allele carriers emerging from drive crosses to escape, persist, and reproduce, thus recolonizing empty space (the resistance allele is usually mostly lost in this process). In contrast, dominant resistance alleles are immediately eliminated when inherited by females, thereby reducing the chance that their carriers can escape to empty space.

### Characteristics of homing suppression drive waves

Based on our preliminary comparison, we suspected that dominant resistance has potential to avoid chasing and accelerate suppression. To gain deeper insight into the underlying dynamics, we simulated the spread of drive waves in a one-dimensional spatial model and estimated wave speed and width, important characteristics for spatial performance^41^. In spatially structured populations, individuals typically spend their lives within a limited local area, mating and reproducing only with nearby neighbors. Consequently, when a gene drive is released into a wild-type population, it generally propagates as a “wave of advance.” Measuring the width and speed of drive wave can provide a useful framework for understanding and comparing the behavior and performance of different drive systems under spatial conditions^41,45^.

Apart from the HSD-dominant and HSD-recessive systems, we also examined the HSD-dominant system with sterile homozygous drive males, which has been experimentally demonstrated^18^. In the simulations, drive carriers were released from the left side of the arena, generating a wave of advance that spread from left to right and displaced the wild-type individuals. All three systems were evaluated under identical parameter settings chosen to represent feasible conditions (drive conversion rate = 0.9, germline resistance rate = 0.1, embryo resistance rate = 0.1, and female drive-carrier fitness = 0.9).

The differences in wave speed among the systems was negligible (Table 1). This is because homing suppression drives will spread mostly based on the drive conversion rate. Suppression effects happening where drive frequency is high won’t substantially affect the leading edge of the wave. However, differences emerged in wave width. The HSD-dominant system exhibited the narrowest width (9.32), followed by the HSD-recessive system (12.08), and then the HSD-dominant system with sterile drive-homozygous males (17.11). Wave width indirectly indicates the capacity of the drive to eliminate the population because rapid elimination will quickly reduce the population when drive frequency is high, leading to a narrower wave. For the HSD-recessive system, when resistance alleles are first generated, they typically occur in resistance/wild-type heterozygotes, which cannot be immediately removed. Thus, the interaction range between drive carriers and wild-type individuals is broader, compared with the HSD-dominant system. For the HSD-dominant system with sterile drive homozygous males, the drive alleles are removed more rapidly in sterile D/D males at the back of wave where the drive is at high frequency, thus slowing population elimination. The eventual removal of wild-type alleles has to rely more heavily on the dominant sterility of resistance alleles, which require a longer time to replace wild-type. Thus, the width of wave is much broader than the other two HSD systems.

**Table 1.**
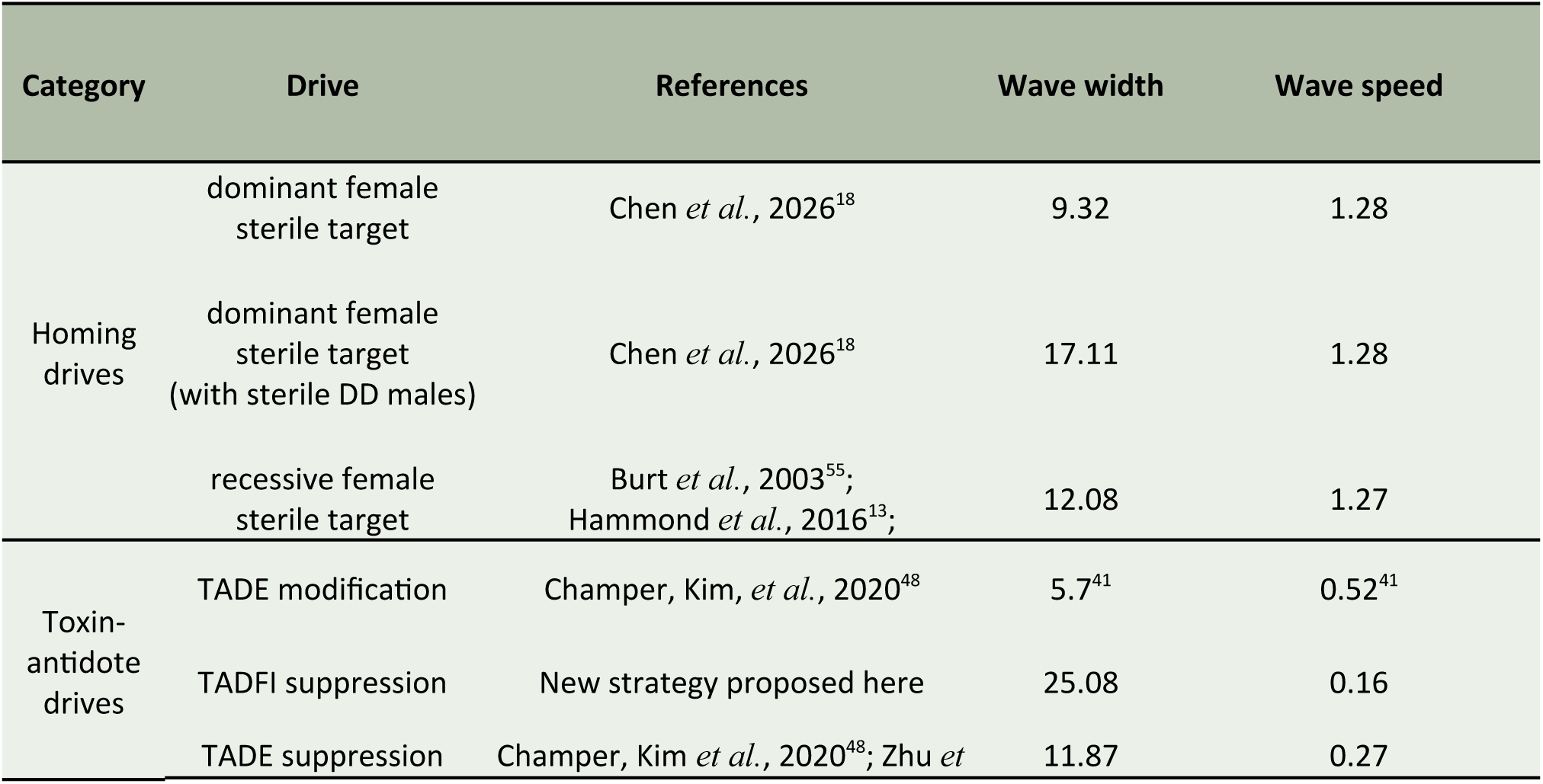
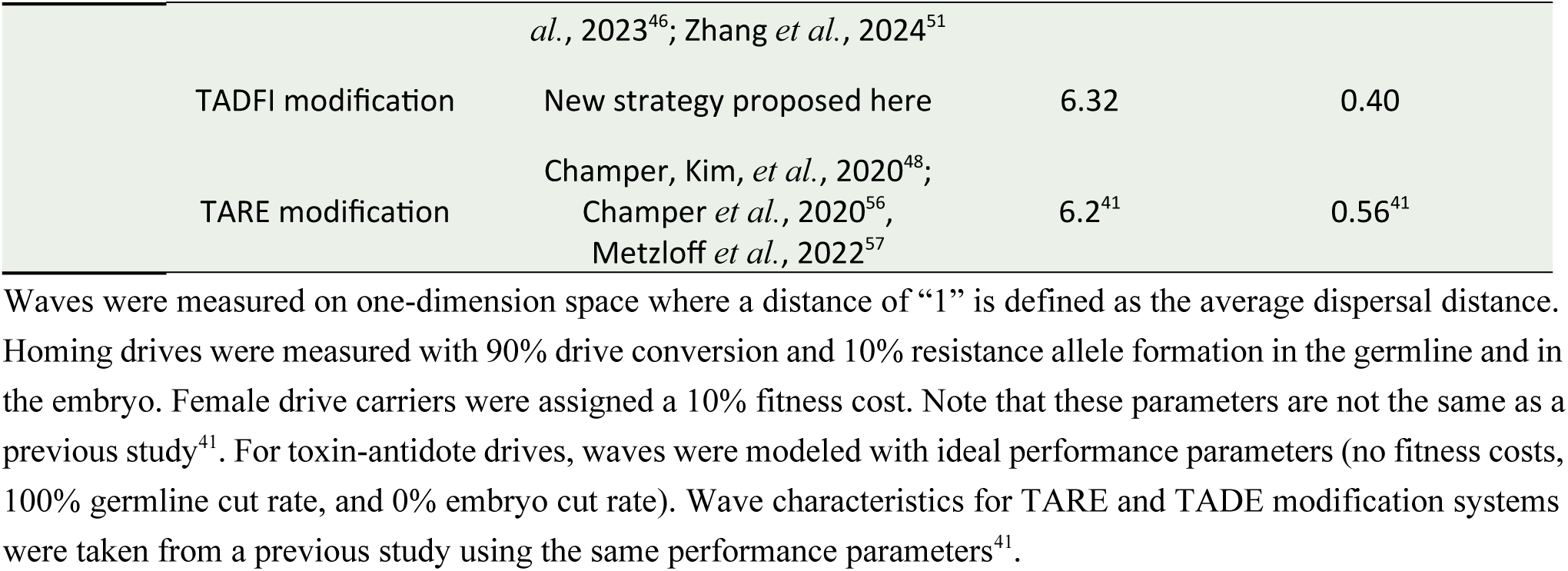
Wave speed and width measurements of drive systems in spatial model.

| Category | Drive | References | Wave width | Wave speed |
| --- | --- | --- | --- | --- |
| Homing drives | dominant female sterile target | Chen *et al.*, 2026<sup>18</sup> | 9.32 | 1.28 |
|  | dominant female sterile target (with sterile DD males) | Chen *et al.*, 2026<sup>18</sup> | 17.11 | 1.28 |
|  | recessive female sterile target | Burt *et al.*, 2003<sup>55</sup>; Hammond *et al.*, 2016<sup>13</sup>; | 12.08 | 1.27 |
| Toxin-antidote drives | TADE modification | Champer, Kim, *et al.*, 2020<sup>48</sup> | 5.7<sup>41</sup> | 0.52<sup>41</sup> |
|  | TADFI suppression | New strategy proposed here | 25.08 | 0.16 |
|  | TADE suppression | Champer, Kim *et al.*, 2020<sup>48</sup>; Zhu *et* | 11.87 | 0.27 |

|  |  |  |  |
| --- | --- | --- | --- |
|  | *al.*, 2023<sup>46</sup>; Zhang *et al.*, 2024<sup>51</sup> |  |  |
| TADFI modification | New strategy proposed here | 6.32 | 0.40 |
| TARE modification | Champer, Kim, *et al.*, 2020<sup>48</sup>;<br>Champer *et al.*, 2020<sup>56</sup>,<br>Metzloff *et al.*, 2022<sup>57</sup> | 6.2<sup>41</sup> | 0.56<sup>41</sup> |

Waves were measured on one-dimension space where a distance of “1” is defined as the average dispersal distance. Homing drives were measured with 90% drive conversion and 10% resistance allele formation in the germline and in the embryo. Female drive carriers were assigned a 10% fitness cost. Note that these parameters are not the same as a previous study<sup>41</sup>. For toxin-antidote drives, waves were modeled with ideal performance parameters (no fitness costs, 100% germline cut rate, and 0% embryo cut rate). Wave characteristics for TARE and TADE modification systems were taken from a previous study using the same performance parameters<sup>41</sup>.

### Performance of suppression drive systems in a spatial model

In two-dimensional models, local suppression by drive provides a chance for wild-type individuals to recolonize, which delays final elimination, or even results in indefinite chasing. Outcomes can be classified as drive loss without chasing, drive loss after chasing, suppression without chasing, suppression after chasing, and long-term persistence of both drive and wild-type alleles, which can involve chasing or equilibrium^40^. To analyze drive outcomes in detail, we analyzed our three systems in the two-dimensional model, varying germline resistance and drive conversion (Fig. 2). We marked the deterministic threshold for sufficient genetic load to achieve suppression, which can help distinguish true chasing and equilibrium status^40^. Parameter sets to the left of this line do not have enough power to eliminate the population and instead remain at long-term equilibrium (though close to the line, they may have a chaotic character similar to chasing due to common local population elimination^40^). Average population size was also collected for better understanding of long-term outcomes. In general, ideal drives could usually eliminate the population, sometimes without chasing. As drive performance worsens, elimination after chasing becomes more common, and eventually long-term chasing or equilibrium. Eventually, the drive loses the ability to even persist, resulting in drive loss without chasing. Other drive loss outcomes are relatively rare.

**Figure 2.**
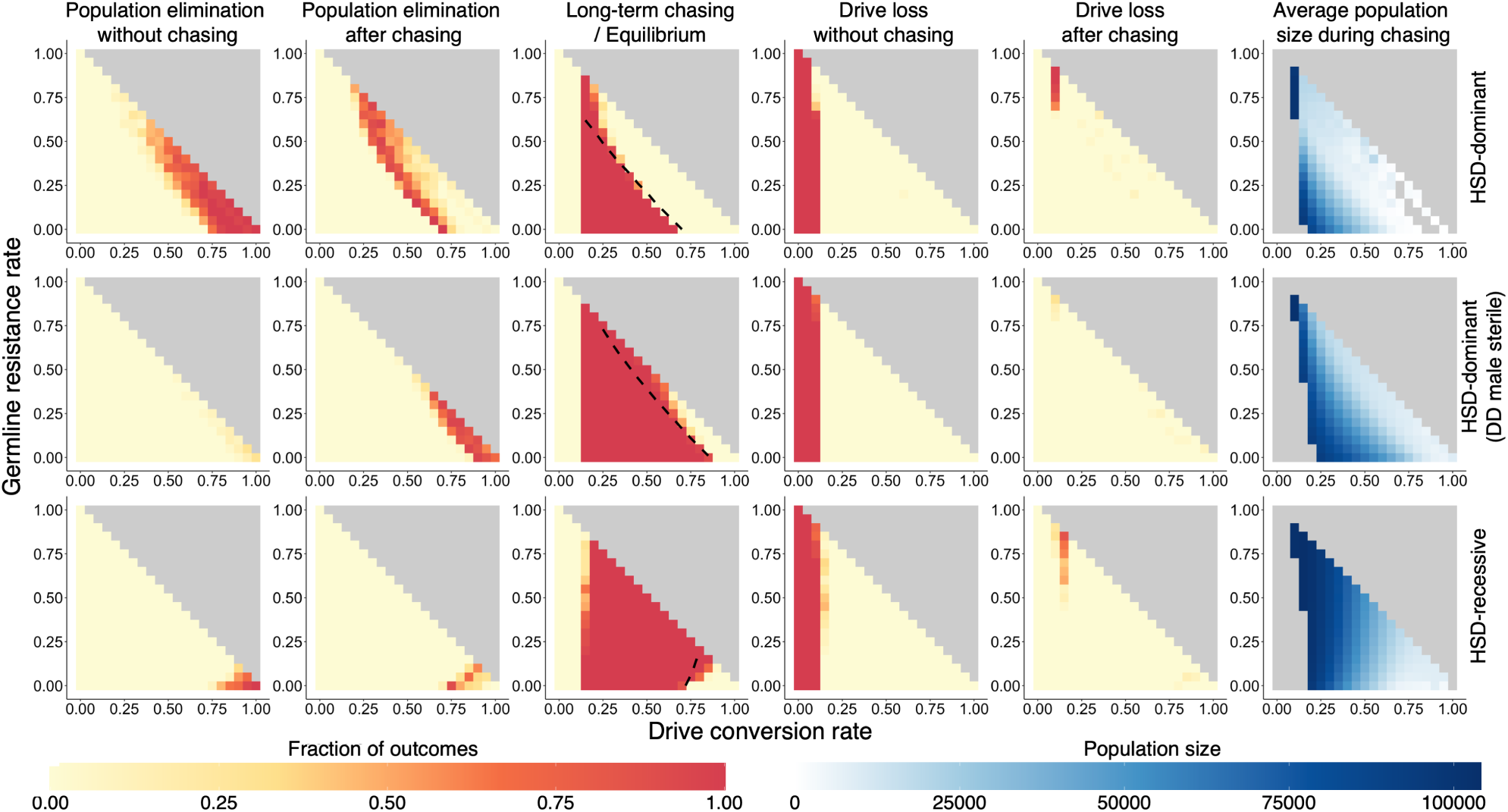
Outcomes of homing suppression drives in a spatial model. Drive heterozygotes representing 10% of the total population were randomly released into a spatial population of 100,000 individuals. Drive carrier female fitness was fixed at 0.8, embryo cutting rate at 0.05, average dispersal at 0.05, and low-density growth at 6. Outcomes are categorized as follows: drive loss without chasing (failure of the drive to establish), drive loss after a period of chasing, suppression without chasing, suppression after chasing, and cases in which both drive and wild-type alleles persisted after 1000 generations (in a state of either chasing or even equilibrium). For simulations involving chasing or equilibrium, the average population size among chasing was recorded during this period. The black dashed line indicates the threshold at which the genetic load is sufficient to achieve suppression in a deterministic model (regions to the right of the line have higher genetic load). Twenty replicate simulations were performed for each parameter set. Other model parameters are at default levels (Supplementary Table 1).

The HSD-dominant system usually has sufficient performance to eliminate the population when the total germline cutting (drive conversion + germline resistance allele formation) rate reaches at least ∼0.8. Suppression can occur even when the drive conversion rate is below 0.5, albeit with a high likelihood of chasing. This is because in this system, resistance allele formation itself exerts suppressive power, partially compensating for the reduced drive conversion. Thus, resistance alleles facilitate rather than hinder drive spread, meaning that the system places relatively modest demands on drive parameters. Drive conversion still needs to be at least 25%, though, to overcome fitness costs with our parameter settings. However, for the HSD-dominant drive with sterile drive homozygous males variant, the parameter range for success is substantially smaller. There is a high probability of chasing occurring, even when the drive has ideal performance parameters, which further illustrates how male homozygous sterility weakens overall performance. However, germline resistance still contributed to drive power, and elimination could still be observed even when the drive conversion rate dropped to 0.65. In the HSD-recessive system, performance was always inferior to the HSD-dominant except when ideal. Compared to the HSD-dominant system with sterile homozygous males the probability of chasing was lower under nearly ideal performance. However, the system is highly sensitive to germline resistance, which hurt performance. Even small increases in germline resistance rates could lead to complete failure, despite high drive conversion (eg. when drive conversion is 0.85). This system thus requires very stringent performance parameters for successful elimination, and was inferior to the HSD-dominant system with sterile drive homozygous males over a substantial portion of the parameter range where total germline cleavage rates were high.

### Tolerance of suppression systems to embryo resistance allele formation

To gain a more comprehensive understanding of the properties of three suppression drive systems, we next evaluated the tolerance to resistance allele formation in the embryo. These are frequently formed due to persistence of maternally deposited Cas9 and gRNA, impeding the spread of the drive if they formed in an embryo that inherits a drive allele^26,30,58,59^. Thus, we tested the drives with varying embryo cut rate (Fig. 3). Interestingly, the HSD-dominant system shows near immunity to embryo cutting in terms out outcomes, achieving elimination while exhibiting only a minimally increased likelihood of chasing even when embryo resistance was 100% (Fig. 3a), though population elimination still occurred more slowly than for simulations with lower embryo resistance. The HSD-dominant system with DD sterile males tolerated embryo cutting rates only below 0.5 before rapidly losing the ability to eliminate the population. The HSD-recessive system showed a steady reduction in elimination success between 0 and 40%.

**Figure 3.**
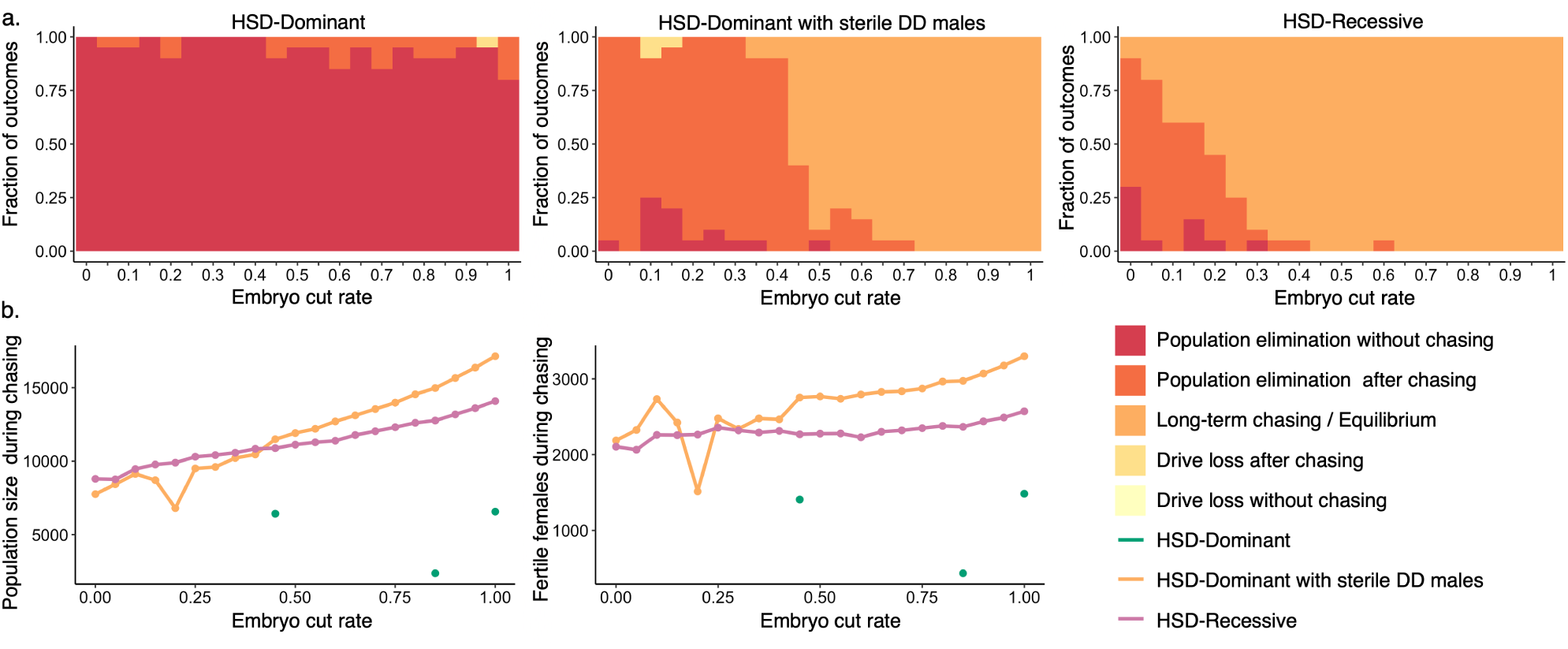
Outcomes of homing drive systems with varying embryo cutting rate in a spatial model. Homing drives (HSD-dominant, HSD-recessive, and HSD-dominant with sterile DD males) were measured with 90% drive conversion, 10% resistance formation in the germline stage, and 20% fitness cost for female drive carriers. Drive heterozygotes representing 10% of the total population were randomly released into a spatial population of 100,000 individuals. **(a)** Outcomes of HSD-dominant, HSD-dominant with DD male sterile, and HSD-recessive systems. Outcomes are categorized as follows: drive loss without chasing (failure of the drive to establish), drive loss after a period of chasing, suppression without chasing, suppression after chasing, and cases in which chasing or equilibrium persisted after 1000 generations. Each parameter set was evaluated with 20 replicate simulations. **(b)** Average population size and average number of fertile females during chasing/equilibrium state, excluding simulations in which the duration of chasing was less than 20 generations. Note that green dots represent the small number of simulations where the HSD-dominant drive had short-term chasing. Note also that the HSD-dominant with sterile drive homozygous males result is affected by stochasticity for the embryo cut rate is low due to the limited duration of chasing in this parameter range.

In addition, we recorded the average population size and the number of fertile females during the chasing phase (Fig. 3b). We excluded simulations in which the duration of chasing was less than 20 generations to reduce the influence of short-term stochastic events. Despite better outcomes when embryo resistance was below 0.6, the HSD-dominant system with DD sterile males retains a slightly larger equilibrium population (both total and fertile female) compared to the HSD-recessive system when embryo cut rate is above 0.4. The advantage conferred by dominant resistance is insufficient to offset the slower spread of the drive caused by the sterility of homozygous drive males.

### Tolerance of suppression systems to functional resistance allele formation

If cleavage is repaired through end-joining, it often produces a mutated target site, giving rise to a resistance allele. In most cases, such resistance alleles disrupt the target gene’s function, usually either by introducing a frameshift or by causing substantial alterations in the amino acid sequence. However, if the target site is less conserved or if an insufficient number of gRNAs is used, functional resistance may arise^13,48,59–61^. Functional resistance both evades drive cleavage and retains gene function, leading to complete failure of suppression drive propagation. Although many studies have shown that the risk of functional resistance can be effectively reduced by increasing the number of gRNAs, it can be a larger problem in chasing or other long-term persistence scenarios because of continued influx of new resistance alleles (which have a small chance of being functional) when drive alleles are together with wild-type^12,20,29,62,63^. Thus, we considered how functional resistance may affect our drives. In our system, dominant-sterile resistance alleles can still override the effect of a functional resistance allele, just like a wild-type allele.

Because functional resistance can have a substantial impact on the outcomes of gene drives, we performed simulations under relatively high-performance drive parameter conditions for all three systems (drive conversion rate = 0.9, germline resistance rate = 0.1, and embryo cutting rate = 0.03). We incorporated a parameter for functional resistance that specified the fraction of total resistance alleles that are functional, varying from 10^-6^ to 10^-3^ (Fig. 4). Because the population capacity is 100,000, a relative functional resistance occurrence rate of 10^-6^ can be regarded as nearly equivalent to the absence of functional resistance in most simulation replicates. The distribution of outcomes across simulations is shown for each drive (Fig. 4). Elimination success rate decreased in all systems as the relative functional resistance rate increased, with failure occurring in nearly all simulations at 10 ^-3^. Both the HSD-dominant with DD male sterility system and the HSD-recessive system showed a high likelihood of chasing, but the HSD-recessive system was moderately more sensitive to functional resistance due to greater chasing duration before elimination. Chasing effectively increased the number of resistance alleles formed before elimination, thus increasing the chance the one will be functional and avoid short-term stochastic elimination. This is why the three drive types suffered similar functional resistance before chasing outcomes due to similar spread properties, but had different rates of functional resistance forming during chasing due to their greatly varying chasing propensity.

**Figure 4.**
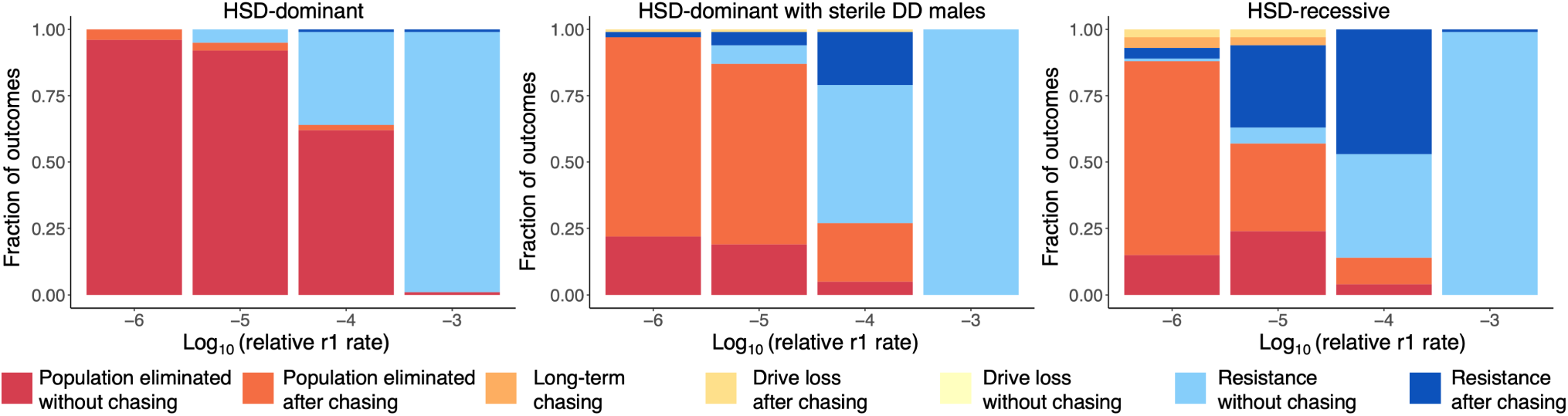
Homing drive systems with varying functional resistance allele formation rate in a spatial model. HSD-dominant, HSD-dominant with sterile DD males, and HSD-recessive drives were measured with 90% drive conversion, 10% resistance generation in the germline stage, 3% embryo cutting rate, and 20% fitness cost for female drive carriers. The relative r1 rate denotes the fraction of functional resistance alleles among all newly generated resistance alleles. Drive heterozygotes representing 10% of the total population were broadly released into a spatial population of 100,000 individuals. Outcomes are categorized as drive loss without chasing (failure of the drive to establish), drive loss after a period of chasing, functional resistance alleles taking over the population without chasing, functional resistance alleles taking over the population after chasing, suppression without chasing, suppression after chasing, and cases in which chasing persisted for 1000 generations. Each parameter set was evaluated with 100 replicate simulations. Other model parameters are at default levels (Supplementary Table 1).

### A threshold-dependent suppression drive based on dominant female sterile alleles

Homing drives are highly efficient, but one of their major problems in some scenarios is the potential to cause global suppression or modification. The number of options for confined suppression are limited. Thus, we propose a threshold-dependent drive based on targeting a female dominant-sterile site, which we term Toxin-Antidote Dominant Female Intersex (TADFI) (Fig. 5a). As a threshold-dependent drive, it is expected to remain confined and thus be suitable for local pest control. If the initial release exceeds the threshold, the drive can establish, spread, and achieve suppression or modification, else the drive allele will fail to propagate and be eliminated. Here, we consider TADFI variants for suppression and modification (Fig. 5a).

**Figure 5.**
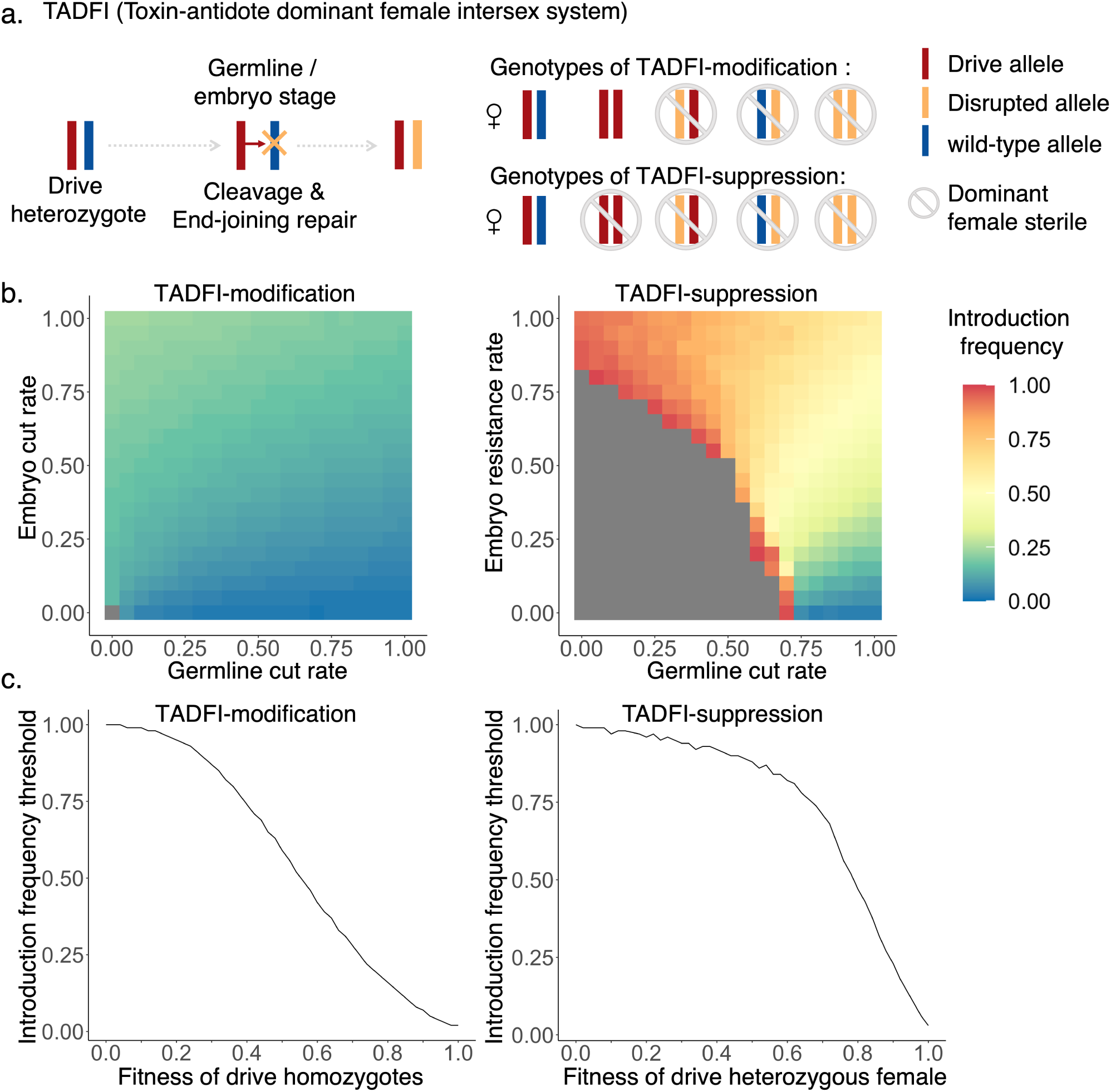
Scheme of TADFI drive system and threshold measurement in panmictic model. **(a)** In the TADFI drive, germline activity disrupts the target gene without drive conversion. Disrupted alleles are dominant female sterile. In the TADFI-modification system, the drive construct contains a female rescue element, so homozygous females are fertile. In the TADFI-suppression system, the drive lacks a rescue copy, so homozygous females are sterile (males remain unaffected by the drive or disrupted alleles). **(b)** Measurement of the introduction frequency threshold for both TADFI modification and suppression drives with no fitness costs under varying embryo and germline cut rates. **(c)** Measurement of the introduction frequency threshold with varying fitness costs. These are specified for drive homozygotes for the modification system (with multiplicative costs per drive allele) and for (heterozygous) drive females for the suppression system. Germline cut rate was set to 100%, while embryo cutting was set to 0%.

Possible design of both TADFI variants can be based on the female-specific splicing site of the *doublesex* gene (Supplementary Fig. 1)^18,28^. TADFI drives are designed to express gRNAs targeting the female-specific splicing site and generate dominant female sterile alleles^28^. Unlike homing systems, the drive allele does not replicate itself through HDR. Instead, its frequency increases by converting wild-type alleles into disrupted (nonfunctional resistance) alleles, which will be subsequently eliminated through dominant female sterility. The drive allele must be designed in a way that avoids causing dominant sterility itself, while also avoiding copying by HDR. For TADFI-modification, the drive should rescue function of both males and females, maintaining the original expression pattern (Fig. 5a). Due to their dominant effect, TADFI is expected to remove disrupted alleles more rapidly than Toxin-Antidote Recessive Embryo (TARE) systems that target essential genes where null alleles are recessive lethal. For TADFI-suppression, the drive must be precisely engineered with a female-specific “rescue” splicing site to block the production of male transcripts in females but not provide functional female transcript, while maintaining normal expression in males (Fig.5a). Thus, homozygous drive females should be sterile (Fig. 5a). Sex-specific splicing elements can be engineered from genes such as *sex-lethal*, *transformer*, or *fruitless*^64–67^. No HDR events should copy the drive in TADFI system, so the design should avoid homology at the cleavage site (though note that a limited amount of drive copying should be acceptable and not change the population dynamics of the system if outweighed by fitness costs). Possible design variants are shown (Supplementary Fig.1). The drive element can be placed in the same site of the gRNA target region, but with truncated homology on either side to wild-type alleles (Supplementary Fig. 1a). The homology regions match to the left of exon 3 and the right of exon 4, which provide >200 bp gap for left side and >1000bp for right side to avoid HDR. Recoded exon 3, a female-specific splicing element, and a terminator (for suppression) / female rescue (for modification) are provided to regulate female and male splicing (Supplementary Fig. 1a). Another design is specifically for suppression (Supplementary Fig. 1b). In this variant, the drive element is inserted in the boundary of exon 2 and the following intron. A terminator will end the original transcript expression in both males and females. At the right of the drive element, a male-specific promotor and recoded CDS region of exon 2 are provided, restarting male transcript expression in only males.

The dynamics of the TADFI systems in a panmictic model is shown in Supplementary Figs. 2-3. In the TADFI modification system, drive and dominant disrupted alleles both increase at first, but then disrupted alleles are rapidly eliminated, and the drive allele takes over the whole population (Supplementary Fig. 2a-b). Note that in our model, we see a suppression effect for the modification drive. This can be attributed to the female dominant effect of the target and sensitive density-dependent growth curve settings and are not necessarily representative of population changes with other models (Supplementary Fig. 2b). For TADFI-suppression drive, when the initial introduction ratio exceeds the threshold, the population has an increasing fraction of drive and disrupted alleles and is ultimately eliminated (Supplementary Fig. 3a-b). However, in both TADFI systems, if the initial introduction frequency is below the threshold, the drive will fail to spread and be eliminated (Supplementary Figs. 2c, 3c).

We further investigated how drive parameters influence the introduction threshold of both TADFI systems (Fig. 5b and c). TADFI suppression drive performs efficiently with high germline cut rate and low embryo cut rate, requiring fewer released individuals to achieve suppression. Small fitness costs or embryo cut rate will increase the threshold. Even with high embryo cut rate, suppression can still be achieved by increasing the release size. Interestingly, embryo cut rate can partially compensate for low germline cut rate, allowing drive spread despite the high release ratio required, though such a scenario is unlikely happen in reality because Cas9 must first be expressed in the germline before it can be maternally deposited. TADFI-modification requires lower initial drive releases, even with low germline cut rate and high embryo cut rate. Fitness is also a key parameter shaping the threshold, so we assessed the threshold under idealized drive parameters but varied fitness (Fig. 5c). For both systems, any fitness cost will significantly increase the introduction threshold frequency.

### Simulation of the TADFI system in a spatial model

Previous studies demonstrated that drive properties, such as confinement, can vary substantially in spatially explicit populations, especially for threshold-dependent confined drive systems^33,38,68–70^. Additionally, suppression drives also introduce additional complexity in spatial scenarios such as chasing. Thus, we explored TADFI’s spread dynamics in a spatially structured population. We first compared the wave of TADFI modification drive to the commonly used TARE system (Table 1), TADFI-modification does not exhibit a clear advantage and in fact spreads somewhat more slowly than both TARE and TADE. The construction of TADFI is also likely more challenging than for the TARE system. We next compared TADFI suppression drive with TADE suppression in the one-dimensional spatial model (Table 1). These two systems have some similarities, as both are designed to target loci with dominant effect to prevent the retention of disrupted alleles in the population^46,48^. TADE suppression drive (in one form) is located in a female fertility gene (thus making the drive recessive sterile) and targets a haplolethal gene, while the drive provide a rescue copy for the haplolethal gene^46,48^. To better compare two systems, we assumed that the TADE drive and target loci are nearby. We observed that the wave width of TADFI is 2.1 times that of TADE suppression, while its spread speed is only 60% as fast (Table 1). In TADE suppression, the dominant effect eliminates non-drive alleles in both males and females, whereas in the TADFI system, the dominant effect occurs only in females, and the effect still allows the individual to compete, potentially reducing resources for other individuals in drive regions. Consequently, the increase in drive frequency is slower in TADFI compared to TADE suppression. However, though the TADE suppression system is stronger, it is difficult to construct^52^. The TADFI system may be easier, though this would still involve overcoming the challenge of achieving rescue in males while avoiding dominant female sterility^18^.

For frequency-dependent drives, chasing can be more detrimental compared to zero-threshold systems^46^. Thus, we further investigated the performance of TADFI suppression drive in continuous space. We first determined the effect of average dispersal distance and low-density growth rate on drive outcomes under ideal drive performance, while fixing the release parameters to ensure drive establishment (a release radius of 0.2 and release ratio of 0.2) (Fig. 4a). The low-density growth rate exerts minimal influence on suppression outcomes unless it falls below 4, in which case large stochastic effects arise that can cause drive loss during chasing. Dispersal has a strong impact on system outcomes. When dispersal is lower than 0.015 (close to the density-interaction distance of 0.01), the drive will be rapidly lost. This is because TADFI cannot form an effective wave advance to invade when the interaction with wild-type population is so restricted, while interaction between drive individuals results in rapid elimination and thus collapse of the wave. As dispersal increased, outcomes shifted to long-term chasing and then elimination after chasing. Elimination without chasing occurred only in occasional cases under very high dispersal. An average dispersal distance of at least ∼0.07 was required to achieve a high probability of successful elimination. The range of long-term chasing is larger compared to TADE-suppression system^46^. Unlike homing drives or other toxin-antidote suppression drives, the phenomena of chasing cannot be avoided for TADFI even when the average dispersal distance is 0.1. This may be because the speed of TADFI system spread is comparable to the recolonization speed of wild-type individuals in empty areas. However, though chasing is common in TADFI system, it can still often catch up and eliminate the population.

Maternal deposition of Cas9 is consistently detrimental to drive-carrying offspring for TADE and TADFI, as it converts their wild-type alleles into disrupted alleles in the embryo stage, leading to lethality or sterility. For both homing and toxin-antidote drives, embryo cutting is an undesirable but often difficult-to-avoid feature. We investigated the outcome of a single widespread TADFI drive release in our spatial model, varying both the embryo cut rate and the release ratio (Fig. 6b). Release ratio refers to the proportion of released individuals in the total population after our release. When the embryo cut rate was less than 0.2, elimination can be achieved easily by increasing the release ratio, despite the possibility of chasing. However, as embryo cut rate continues to increase, the release ratio must exceed 0.8 for successful population elimination. Otherwise, the drive will be lost, either rapidly due to failure to exceed the drive threshold or after a period of chasing, where the drive will be unable to form an effective wave of advance. With a high release level, the population can be eliminated without chasing, even with very high embryo cutting. This is because embryo cutting does not affect the spreading efficiency of males. Instead, it only diminishes the transmission capability through the daughters of drive-carrying females. Therefore, by increasing the release size to boost the initial drive frequency, rapid population elimination can still be achieved with a single release.

**Figure 6.**
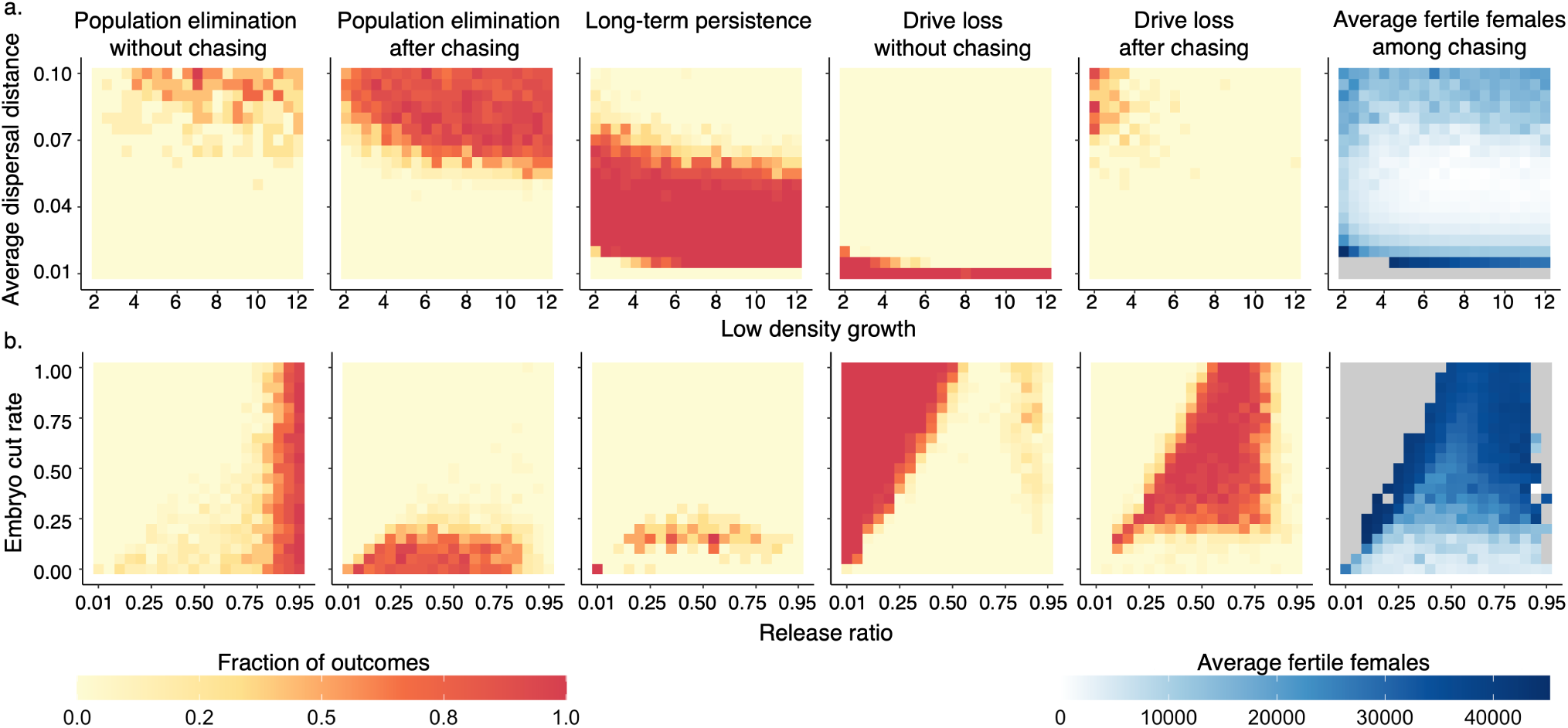
Outcomes of the TADFI-suppression system in a spatial model. Drive heterozygotes were released into a population of 100,000. Germline cut rate was 100%, and no fitness costs were considered. Outcomes are categorized as follows: drive loss without chasing (failure of the drive to establish), drive loss after a period of chasing, suppression without chasing, suppression after chasing, and cases in which both drive and wild-type alleles persisted for 1000 generations (either chasing, equilibrium, or slow drive growth). Each parameter set was evaluated with 20 replicate simulations. **(a)** Outcomes under different low-density growth and average dispersal distance . Fixed parameters included a release ratio = 0.2 and 0% embryo cut rate. Drive heterozygotes were released in a central circle of radius 0.2. **(b)** Outcomes under varying embryo cut rates and release ratios, with fixed ecological parameters (average dispersal = 0.08; low-density growth rate = 6). Drive heterozygotes were randomly released. Other model parameters are at default levels (Supplementary Table 1).

## Discussion

In our spatial model, mating occurs within a limited distance, and competition is local and density-dependent. Escaped wild-type individuals have a higher chance of recolonizing empty areas, and thus, the conditions required for population elimination become more stringent. Even good-performing drives in the panmictic model may fail to achieve suppression in the spatial model (Fig. 1). Previous studies suggested that targeting dominant sites can enhance the suppressive power of homing drives, but it remains necessary to test whether dominant fertility targets retain this advantage under more realistic release scenarios^18,28^. The commonly used HSD-recessive system (Homing suppression drive targeting recessive female fertility gene) was highly vulnerable to resistance alleles. Suppression was achieved only when the drive conversion rate was at least 75%. High germline resistance substantially increased the probability of chasing and impeded population elimination. In contrast, we found that the HSD-dominant system can achieve population suppression even with low drive conversion rate, when the total germline cut rate is sufficiently high (>75%) (Fig. 2). This effect can be attributed to dominant female sterility, which prevents escaped resistance allele carriers from recolonizing the population. This comparison highlights the advantage of targeting dominant sites. They not only increase suppressive power, but also allow avoiding chasing over a large parameter range. Even with inferior performance parameters that produce equal genetic load to HSD-recessive systems, they can avoid long-term chasing when HSD-recessive systems remain vulnerable.

However, the HSD-dominant system is difficult to construct^18^. The dominant effect of disrupting *doublesex* is achieved by incorrect splicing in females, but the drive element at the splicing site also tends to interfere with normal splicing in males. Drive homozygous males lose fertility, which reduces overall drive effectiveness. ’Chasing’ is more prevalent in HSD-dominant systems when drive homozygous males are sterile (Fig. 2). Although such systems do not show a clear advantage in avoiding chasing compared to standard suppression systems, they still enable more effective population suppression under high germline cutting rates, even when the drive conversion rate is modest. For example, the population could be eliminated after chasing when the drive conversion rate was only 65% if remaining wild-type alleles were converted to resistance alleles (Fig. 2). Thus, even this less effective system has advantages over an important part of the parameter range.

Key limitations of suppression drives are that embryo cutting caused by maternally deposited Cas9 remains difficult to avoid in many species, and functional resistance can completely stop a suppression drive. We examined the effects of functional resistance and embryo resistance on the performance of our homing suppression drive systems (Fig. 3 and Fig. 4). The HSD-dominant system shows a clear advantage in both situations. It is slowed by embryo resistance but retains high suppressive power. Functional resistance alleles are less problematic because chasing can be avoided. For HSD-dominant system with sterile males and HSD-recessive system, embryo cutting rate increases the probability of chasing, though it has less of an impact compared to the standard suppression drive. It has somewhat improved tolerance to functional resistance due to shorter chasing duration before population elimination usually occurs.

Mutations that are dominant female-sterile are comparatively difficult to identify. Based on previous studies, dominant female-sterile effects within *doublesex* targets have been reported in *Drosophila*, *Anopheles*, and *Culex*^16,18,28,71^. As *doublesex* is highly conserved in dipterans, target sites with similar pattern might be achievable in related species. Resolving the issue of homozygous drive males for *doublesex* remains a priority. Our previous study failed to prevent incorrect splicing in homozygous drive males in *Drosophila melanogaster*^18^, and a similar splicing issue was seen in *Anopheles stephensi*^15^. Future studies could consider sex-specific splicing elements to rectify this issue from *sex-lethal*, *transformer*, and *fruitless* genes^64–67^, but careful attention should be paid to whether the expression timing of regulatory proteins for sex-specific splicing aligns with the developmental window during which *dsx* is required for sex determination. Alternatively, other dominant female-specific genes could be explored as potential target sites. For example, the female-specific dominant flight-related gene *myo-fem* in *Aedes* can be a possible target candidate^72^. Recently published transcriptome data can facilitate screening additional candidate genes^73–76^.

The HSD-dominant system is undoubtedly the most effective of the suppression systems we considered, but the construction remains challenging. The HSD-recessive system should have nearly equal performance in species that can achieve exceptionally high drive conversion rates, such as *Anopheles*. HSD-recessive drive allows a broader range of target genes and is relatively easier to construct. The HSD-dominant system with sterile males could also be constructed in a straightforward manner if a suitable target site is found at *dsx*. It exhibits high tolerance to low conversion efficiency if total germline cleavage is high, and it is more tolerant to high embryo cutting than HSD-recessive. It may be suitable for species where high HDR rates are hard to achieve.

Many potential real-world gene drive scenarios require local and controllable strategies. Confined population suppression is often difficult to achieve, relying on difficult-to-construct TADE suppression systems^46,48,49,51,52^ or tethered gene drives that require excellent homing drive performance^57,77,78^. We thus proposed a toxin-antidote gene drive system based on female-specific dominant genes, termed TADFI. This system includes both threshold-dependent modification and suppression variants (Fig. 5). The TADFI-modification system does not offer a clear advantage over TARE (Table 1) and is more challenging to construct, but the suppression system provides potentially advantageous capabilities compared to other designs. In general, TADFI performs well, gaining many of the usual advantages of CRISPR toxin-antidote drives, essentially functioning as a somewhat slower version of TADE. These advantages include lack of need for homology-directed repair copying of drive and a low introduction threshold when performance is good.

We conducted a comparison between the TADFI systems and other toxin-antidote systems in spatial models. The wave of TADFI suppression is slower compared to TADE suppression (Table 1) because disrupted alleles are only eliminated when in females. TADFI becomes less tolerant to embryo resistance in the spatial model because embryo resistance renders drive-carrying females sterile. The propagation of the drive cannot overcome recolonization of wild-type, leading to chasing. We also found that TADFI-suppression has a high likelihood of chasing even with good performance parameters. When the dispersal is relatively high, though, it is less vulnerable to chasing, allowing more successful population elimination outcomes. This suggests that the TADFI system may be particularly sensitive to species ecology. Thus, for accurate predictions, it is essential to make a specific-species model with accurate ecological information.

For the experimental construction of TADFI-suppression, we show a possible design based on *doublesex* (Supplementary Fig. 1). The main challenge is how to maintain sex-specific splicing in the drive allele. The rescue element design logic is similar to that of the HSD-dominant system (see above), and future studies could apply endogenous sex-specific splicing elements from the target species.

Overall, we evaluated the performance of the suppression gene drives with dominant female sterile resistance alleles system in complex spatial environments from multiple perspectives. The results highlight the advantages of using a dominant sterile target site, providing theoretical support and guidance for the design of future homing drives. In addition, we proposed and tested the performance of the TADFI system, offering a new confined suppression option for genetic control strategies.

## Acknowledgements

This study was supported by the Center for Life Sciences and the National Natural Science Foundation of China (grants 32270672 and W2432018). The cluster-based data collection was assisted by High-Performance Computing Platform of the Center for Life Science at Peking University.

---

**Supplementary Figure 1.**
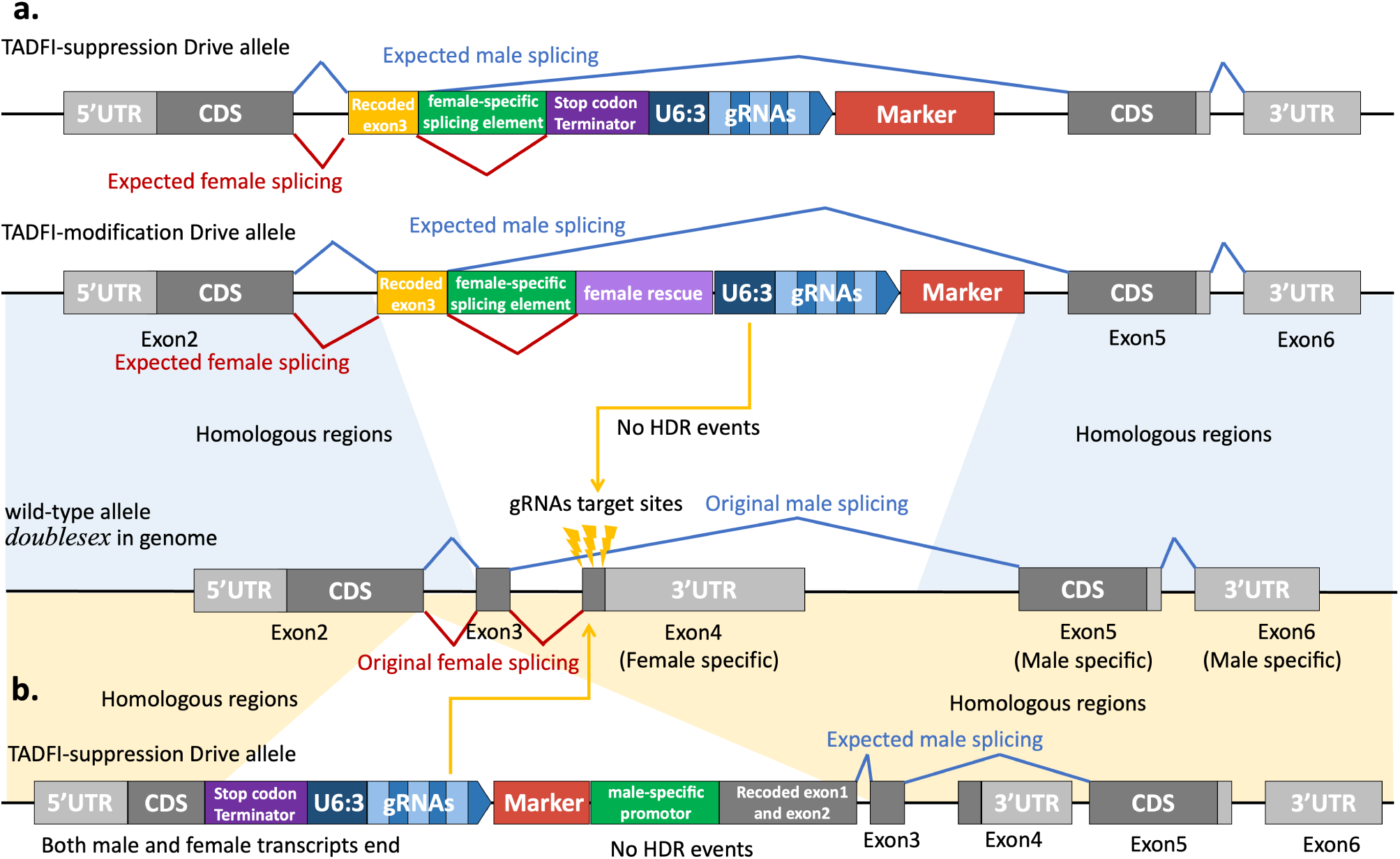
Possible designs of TADFI systems targeting *doublesex*. gRNAs are designed to target the dominant female sterile site (the boundary of intron 3 and exon 4) of *doublesex*. Blue lines show expected/original male splicing, while red lines show expected/original female splicing. (a) The drive element is placed at the same region of target site. Given the large gaps (>200 bp on the left and >1000 bp on the right) between the homologous regions and the gRNA cleavage sites, no HDR events are expected. In TADFI-suppression system, the rescue element only helps with sex-specific splicing. In TADFI-modification system, the rescue element must restore both proper splicing and the function of the female transcript. (b) The drive element is placed in exon 2, ∼24,000bp away from target sites. Both original transcripts end by inserted terminator. A male-specific promotor restarts male transcript expression. Sex-specific splicing elements can be engineered from genes such as *sxl2*, *tra*, or *fru*.

**Supplementary Figure 2.**
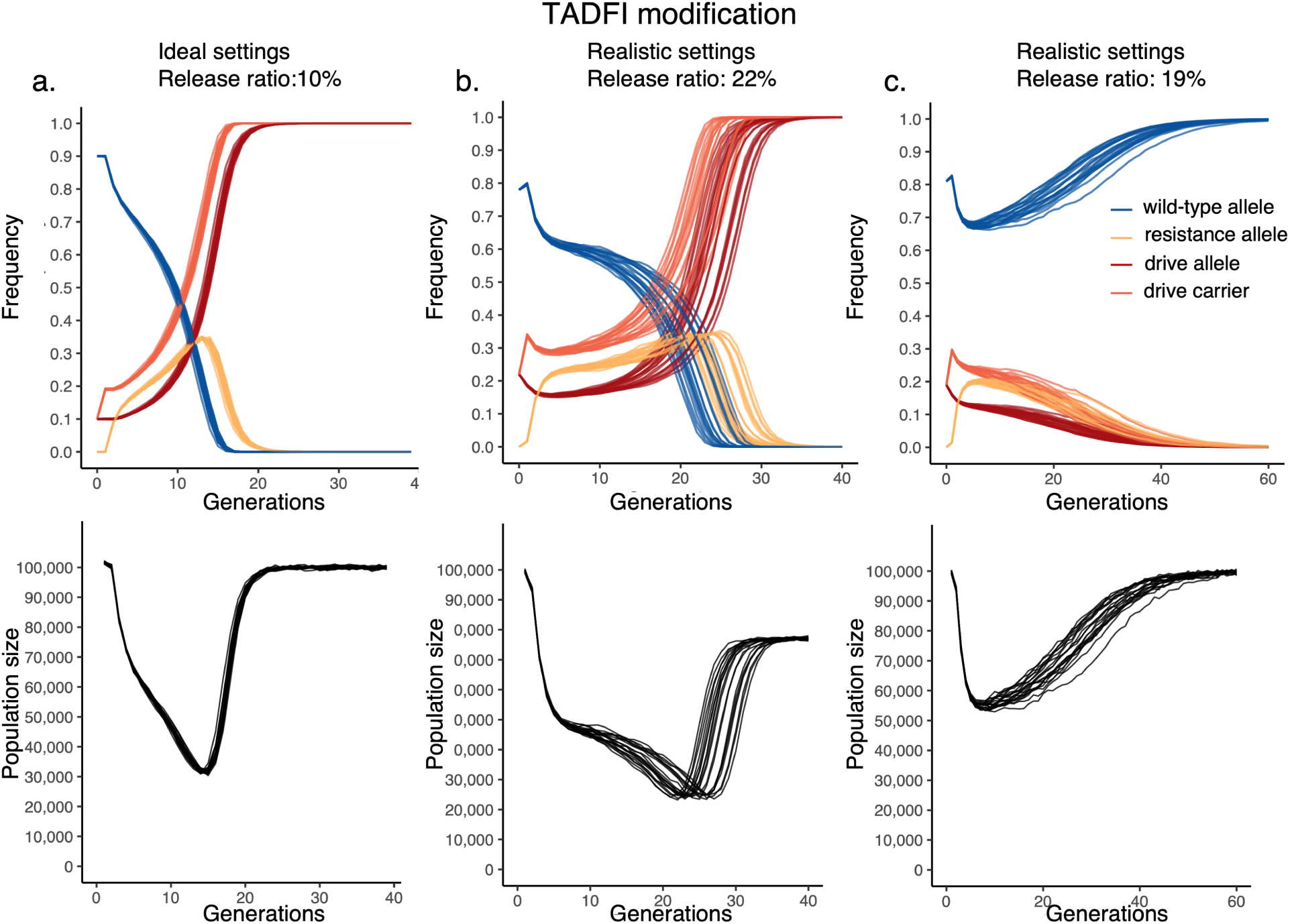
Population dynamics of TADFI modification drive in a panmictic model. The initial population consists of 100,000 individuals. 20 simulations were recorded for each test. **(a)** TADFI modification drive under ideal parameters. Germline cut rate was 100%, with no embryo cutting and no fitness cost imposed on drive alleles. Homozygous drive individuals were released at a frequency of 10%. (b-c) TADFI modification drive under realistic parameters. Germline cut rate was 90%, with 10% embryo cutting and 10% fitness cost imposed on each drive allele. Heterozygous drive individuals were introduced at a frequency of **(b)** 22% or **(c)** 19%.

**Supplementary Figure 3.**
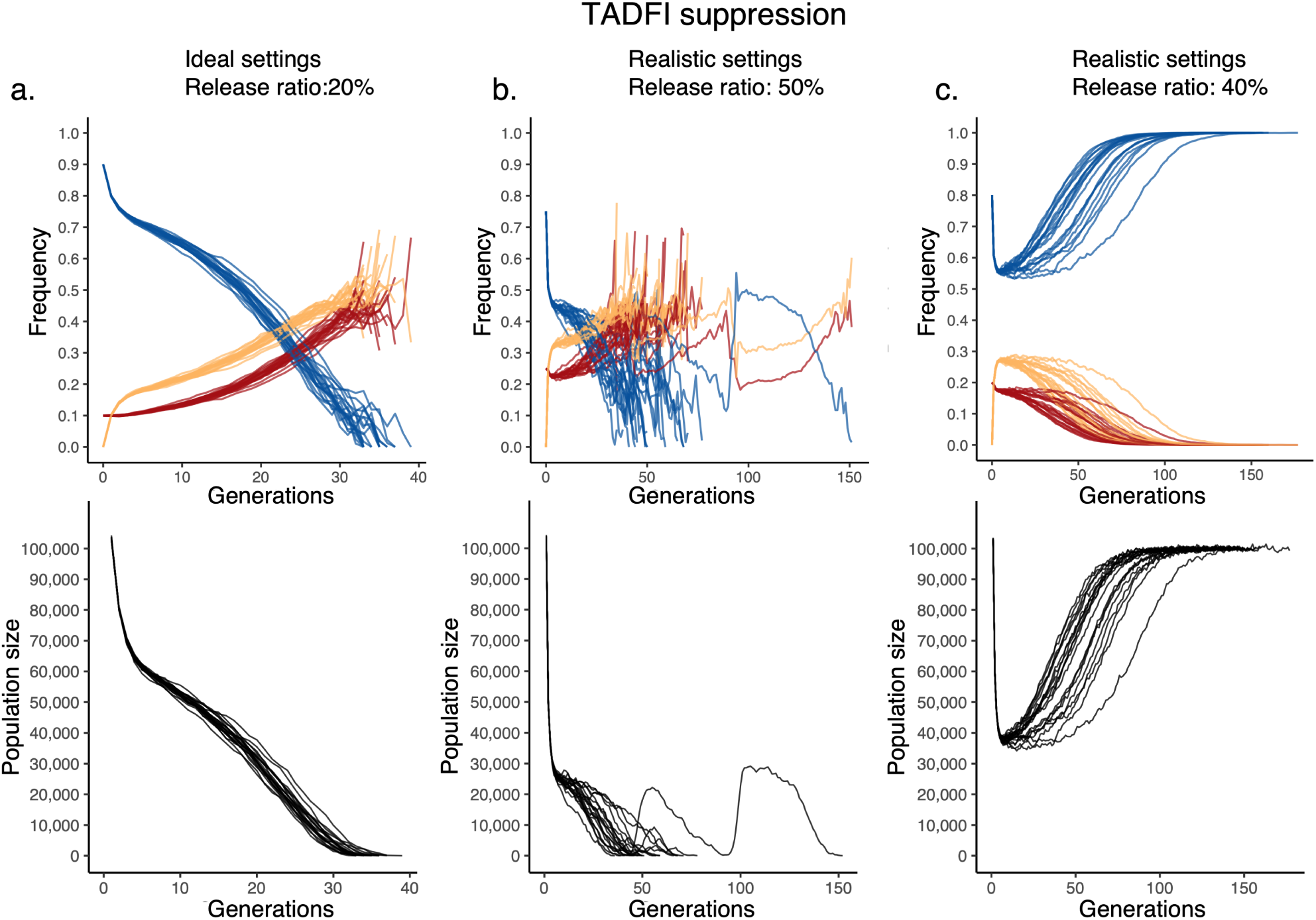
Population dynamics of TADFI suppression drive in a panmictic model. The initial population consists of 100,000 individuals. 20 simulations were recorded for each test. **(a)** TADFI suppression under ideal parameters. Germline cut rate was 100%, with no embryo cutting and no fitness cost imposed on drive females. Heterozygous drive individuals were released at a frequency of 20%. **(b-c)** TADFI suppression under realistic parameters. Germline cut rate was 90%, with 10% embryo cutting and 10% fitness cost imposed on drive females. Heterozygous drive individuals were introduced a frequency of **(b)** 50% or **(c)** 40%.

**Supplementary Table 1.**
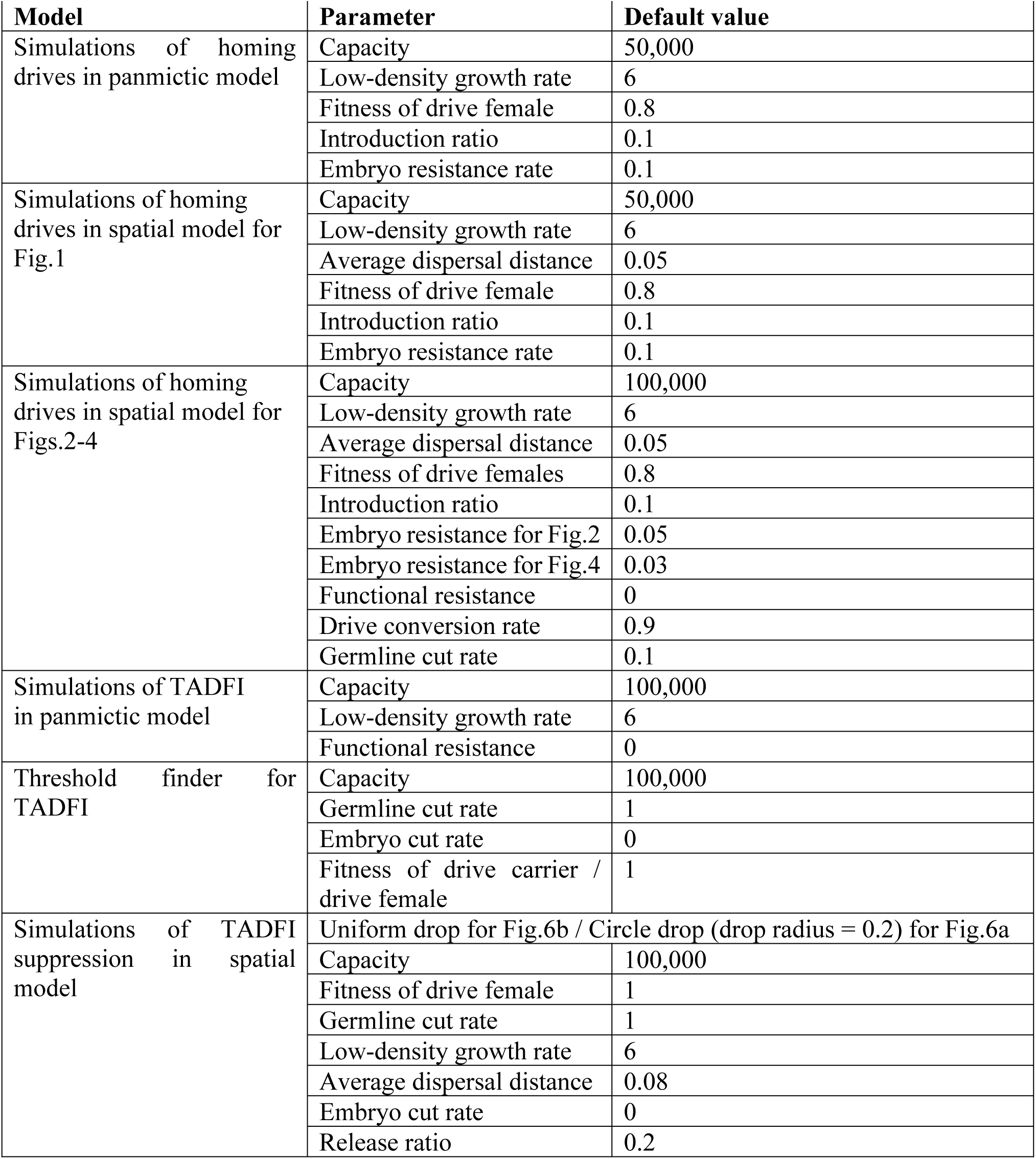
Default parameter settings

## Notes

https://github.com/chenwz22/TADFI_supplement/

### Competing Interest Statement

The authors have declared no competing interest.

